# Sensory attenuation is related to dopamine dose in Parkinson’s disease

**DOI:** 10.1101/221317

**Authors:** Noham Wolpe, Jiaxiang Zhang, Cristina Nombela, James N Ingram, Daniel M Wolpert, Cam-CAN, James B. Rowe

## Abstract

Abnormal initiation and control of voluntary movements are among the principal manifestations of Parkinson’s disease (PD). However, the processes underlying these abnormalities and their potential remediation by dopamine treatment remain poorly understood. Normally, movements depend on the integration of sensory information with the predicted consequences of action. This integration leads to a suppression in the intensity of predicted sensations, and increases the relative salience of unexpected stimuli to facilitate the control of movements. We examined this integration process and its relation to dopamine in PD, by measuring sensorimotor attenuation – the reduction in the perceived intensity of predicted sensations from self-generated versus external actions. Patients with idiopathic PD (n=18) and population-derived controls (n=175) matched a set of target forces applied to their left index finger by a torque motor. To match the force, participants either pressed with their right index finger (‘Direct’ condition) or used a linear potentiometer that controlled a motor (‘Slider’ condition). We found that despite changes in sensitivity to different forces, overall sensory attenuation did not differ between medicated PD patients and controls. Importantly, the degree of attenuation was negatively related to PD motor severity but positively related to individual patient dopamine dose, as measured by levodopa dose equivalency. The results suggest that dopamine could regulate the integration of sensorimotor prediction with sensory information to facilitate the control of voluntary movements.

## INTRODUCTION

A key manifestation of Parkinson’s disease (PD) is bradykinesia – that is, patients have marked difficulties in planning, initiating and executing voluntary movements (Jankovic, 2008). This principal abnormality in motor control has been shown to correlate well with dopamine disruption in patients (Vingerhoets *et al*, 1997), however, the exact mechanism remains poorly understood. Normal motor control depends on the integration of peripheral sensory information with predictions arising from internal models of action. The integration is dependent on the relative ‘precision’ of sensory information and predictions (Franklin and Wolpert, 2011), such that in an uncertain environment, for example, people’s movements rely more on prediction (Körding and Wolpert, 2004). Dopamine has been suggested to play a central role in regulating the precision of sensory information relative to predictions (Friston *et al*, 2012). Striatal dopamine deficit, which is a hallmark pathological feature in PD, is therefore expected to lead to reduced reliance on sensory information (Vilares and Kording, 2017; Wolpe *et al*, 2015) and reduced sensory sensitivity (Konczak *et al*, 2012). However, PD also substantially increases the relative reliance on sensory information; for example, PD patients are more dependent on sensory cues for initiating movements (Morris *et al*, 1996), and the withdrawal of visual feedback impairs patients more than healthy individuals in terms of both movement speed and accuracy (Klockgether and Dichgans, 1994). Here, we examine the integration between sensory signals and predictions in PD, through sensorimotor attenuation.

Sensorimotor attenuation is the reduction in the perceived intensity of stimuli generated by one’s actions, compared to externally generated stimuli. It reflects the suppression of predicted sensory consequences from perception (Blakemore *et al*, 1998). Intact precision of sensorimotor predictions are thought to be required for increasing the salience of external events, to facilitate the rapid initiation (Brown *et al*, 2013) and correction of movements to unpredicted events (Franklin and Wolpert, 2011). In schizophrenia, for example, reduced sensory attenuation and ‘exaggerated’ increase in reliance on sensory information have been suggested to contribute to deficits in distinguishing between self-caused and external stimuli (Shergill *et al*, 2005). Deficits in the integration of prior prediction and sensory information, as reflected in sensory attenuation, can therefore shed light on the mechanism of neurological and psychiatric disorders (Pareés *et al*, 2014; Wolpe and Rowe, 2014).

Sensory attenuation can be quantified by the force matching task. In the force matching task (Shergill *et al*, 2003), 98% of adults show attenuation (Wolpe *et al*, 2016), applying a larger force when matching an external force directly with their hand (‘Direct’ condition). In contrast, people tend to be accurate when matching the force indirectly with a linear potentiometer that controls a motor (Shergill *et al*, 2003). The overcompensation of forces in the Direct condition is associated with the integrity of a fronto-striatal network (Wolpe *et al*, 2016) that is strongly affected by dopamine deficits in PD (e.g. Lewis *et al*, 2003).

We tested patients with idiopathic PD on a force matching task to measure sensory attenuation. Patient measures were compared to normative data from a large epidemiological control cohort (Shafto *et al*, 2014). Patients were tested while ‘on’, after taking their regular dopaminergic medication, and we took advantage of the variability in disease severity and medication to examine between-subject differences in attenuation in relation to motor severity of PD and levodopa dose equivalency. Our principal hypothesis was that PD patients would show changes in attenuation that would reflect abnormal integration of sensory information with sensorimotor prediction (Macerollo *et al*, 2016). We also hypothesised that dopamine medication dose would be related to an alleviation of the effect of PD on attenuation.

## MATERIALS AND METHODS

### Participants

Eighteen patients (12 men; aged 48-81 years, mean: 67; SD: 10) were recruited from the John van Geest Centre for Brain Repair, Parkinson’s disease research clinic. Patients met clinical diagnostic criteria of idiopathic PD, according to the UK PD brain bank criteria (Hughes *et al*, 1992), and were in the mild to moderate stages of disease [Hoehn and Yahr stages 1 to 3] (Hoehn and Yahr, 1967). Normative, population-derived controls were drawn from the Cambridge Centre for Ageing and Neuroscience (<u>https://camcan-archive.mrc-cbu.cam.ac.uk/dataaccess/</u>). Control subjects were selected from the data repository by age, such that all subjects within the patient age range were included in the study (n = 175, 89 men, mean age: 65, SD: 10). All subjects gave written informed consent. The study was approved by the Cambridge Research Ethics Committee.

Assessment of motor and cognitive features in patients was performed at the beginning of the testing session. The severity of motor features was assessed with the Unified Parkinson’s Disease Rating Scale, motor subscale III (Fahn and Elton, 1987). Cognition was assessed with the Mini-Mental State Examination (Folstein *et al*, 1975) and Addenbrooke’s Cognitive Examination Revised (Mioshi *et al*, 2006), excluding patients with ACE-R score below 84/100 (Reyes *et al*, 2009). Patients were tested in the morning after taking their medication as normal. The time interval between levodopa self-administration and testing varied between one and three hours, such that all patients were in a relative ‘on’ state at the time they were assessed. Levodopa dose equivalency (LDE) was computed according to Tomlinson et al. (2010). Clinical data were collected and scored independently and blind to the behavioural results.

### Force matching task procedures and analyses

On each trial of the Force Matching Task (Shergill *et al*, 2003), a lever attached to a torque motor applied the target force for 2.5 s to the left index finger (Fig. 1). The target force was pseudo-randomly selected from the set: 1.0, 1.5, 2.0 and 2.5 Newton (N): each target force was presented once within a cycle of four trials. At the end of the presentation period, the force was removed, and participants used their right index finger to match the force they had just sensed on their left finger (matching period 4.5 sec long). Premature (response during the presentation period) or late (> 1 s) responses led to a warning “too fast” or “too slow” on the computer screen, and the trial was repeated. Because PD patients have altered force output [e.g. increased force irregularities and time to peak (Stelmach *et al*, 1989)], the matched force was calculated as the mean force measured within a 500 ms time window that was selected on a trial-by-trial basis. A sliding window was used to identify the 500 ms interval that had the minimum force variability. This procedure was implemented in both patient and control data.

**Figure 1.**
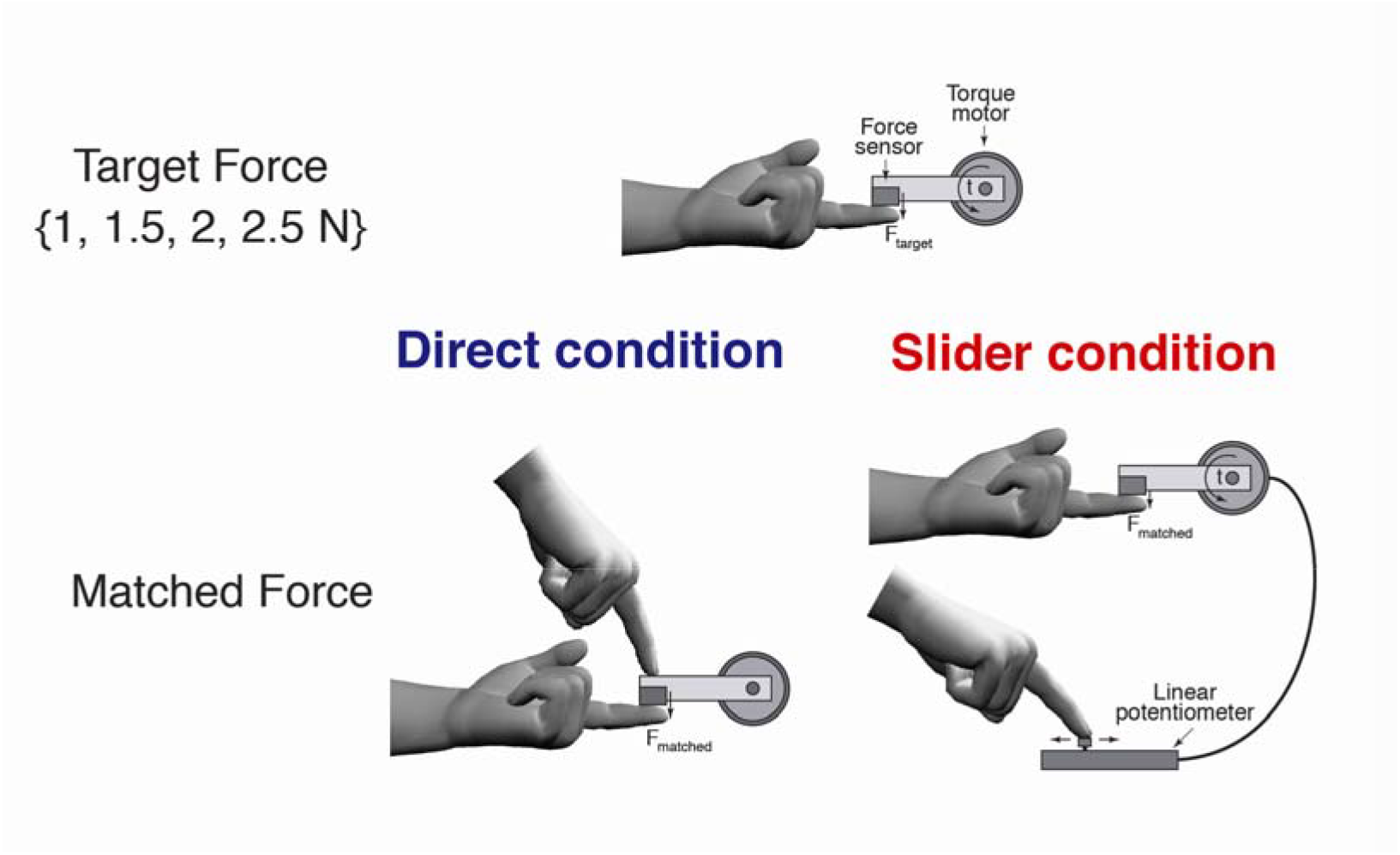
Force matching task illustration. Illustration of the force matching task. In each trial, a torque motor pseudorandomly applied one of four force levels (target force) through a lever to the participants’ left index finger. Participants were asked to match the force they had just sensed (matched force) either by pressing the lever with their right index finger (‘Direct’ condition); or by sliding a linear potentiometer which controlled the torque motor (‘Slider’ condition).

There were two conditions. In the Direct condition, participants matched the target force by pressing with their right index finger directly on top of the lever, mechanically transmitting the force to the left finger. In the Slider condition, participants matched the force with their right index finger by moving a slider (a linear potentiometer) which controlled the torque motor. A force sensor at the end of the lever measured both the target and matched forces applied to the left finger. All participants performed both the Direct and Slider conditions, in a counterbalanced order. For each condition, an initial familiarisation phase of eight trials (two cycles of the four target forces) was performed. The main experiment for each condition consisted of 32 trials.

For each condition, mean force overcompensation was calculated as the average difference between the matched and the target force on each trial across the four target force levels. Positive mean overcompensation reflected an overall attenuation of sensations. To examine attenuation as a function of force levels, the intercept and slope from a linear regression of matched versus target force was calculated for each participant and condition. All statistical tests of behavioural data were performed with two-tailed tests, implemented with ‘R’ software (R Core Team, 2013). Given the unequal sample size (Mann and Whitney, 1947), non-parametric Mann-Whitney test was used for the comparison between groups. After assessing the normality of errors using Kolmogorov-Smirnoff test, within PD group tests were performed with parametric tests. To examine the relationship between force matching and clinical variables in the PD group, multiple regression analyses were conducted with attenuation measures as the dependent variables. The independent variables were disease severity and patient LDE. Covariates of no interest included the variables where patients differed from controls (see Results). All variables were centred and Z-scaled before being entered into the model.

## RESULTS

### Participant demographics

Patient clinical information is summarised in Table 1. Patients (n=18) and controls (n=175) were not different in terms of age (Mann-Whitney test; *Z* = 0.92, *p* = 0.36) and gender *χ^2^* = 1.64, *p* = 0.2).

**Table 1.**
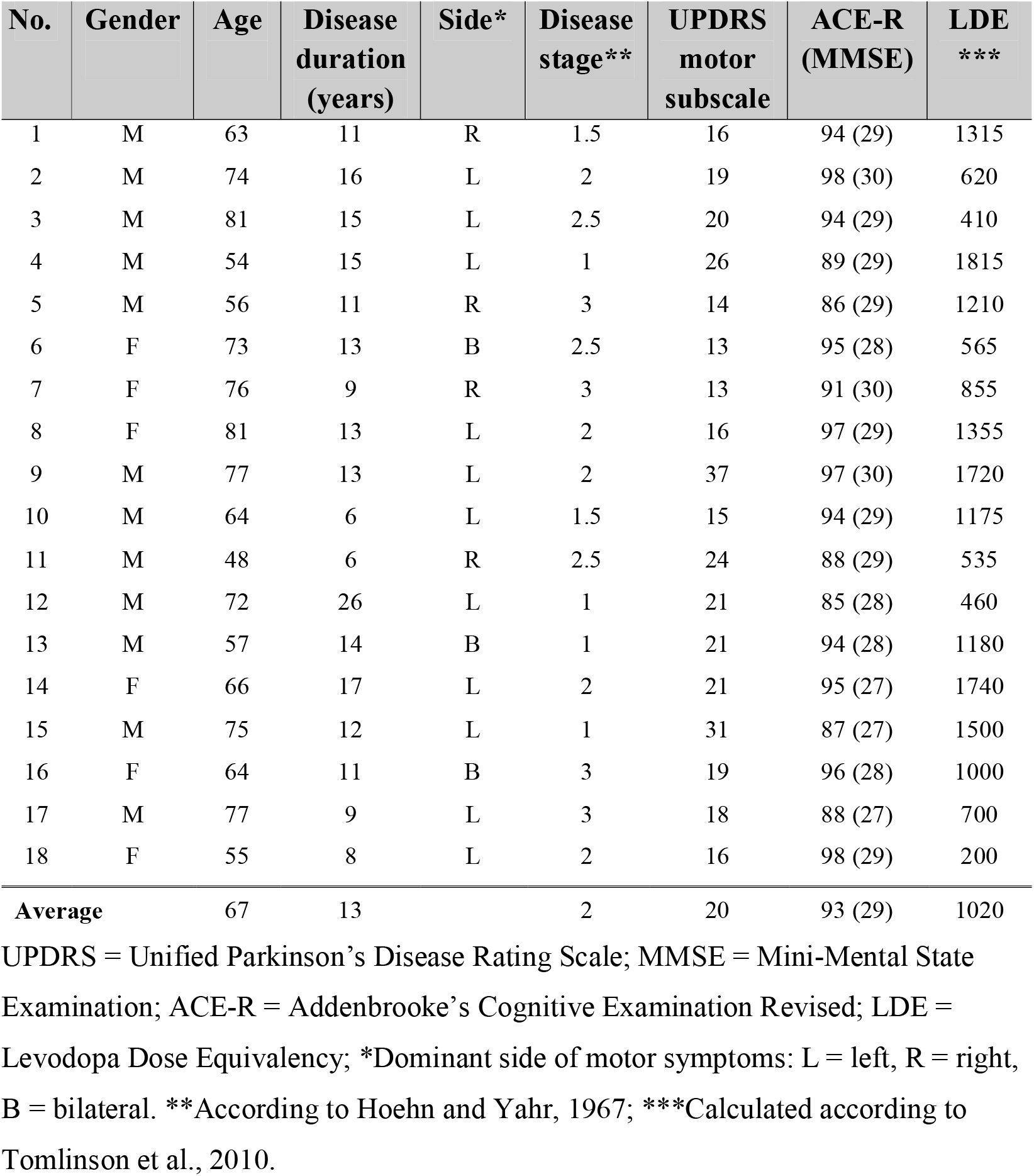
Summary of patient clinical information.

### Sensory attenuation in PD

In order to calculate the matched force for each participant and trial, we calculated the mean force during a 500 ms interval that had the minimum force variability, found with a sliding window. There was no significant difference between initiation times of force matching in patients and controls (Mann-Whitney test; *Z* = −1.58, *p* = 0.11). Importantly the mean time at which patients and controls stabilised the force after initiation was not different (2.67 s versus 2.73 s in patients and controls; Mann-Whitney test; *Z* = −0.37, *p* = 0.71). In the Slider condition, initiation times of force matching were similar across groups (*Z* = 1.39, *p* = 0.16), and patients stabilised the forces slower than controls (2.61 s versus 2.36 s in patients and controls; Mann-Whitney test; *Z* = 2.78, *p* < 0.01). The distributions of calculated mean overcompensation (mean difference between matched and target force) for both conditions in patients were not different from normal distribution (one-sample Kolmogorov-Smirnov test; Direct: *D* = 0.228, *p* = 0.26, Slider: *D* = 0.137, *p* = 0.845).

All patients showed overall sensory attenuation, as indicated by a positive mean overcompensation in the Direct condition (Fig. 2A). Mean Direct force overcompensation was greater than zero (*t_17_* = 6.94, *p* < 0.001), at 1.51 N (*SD* = 0.92 N). Patients were more accurate when matching the force in the Slider condition (Fig. 2A), with smaller force overcompensation than in the Direct condition (*t_17_* = −5.93, *p* < 0.001). Mean Slider force overcompensation was −0.01 N (*SD* = 0.36 N), and was not significantly different from zero (*t_17_* = −0.12, *p* = 0.91). These results confirm that sensory attenuation is as robust in PD patients as in the general population (Wolpe *et al*, 2016).

**Figure 2.**
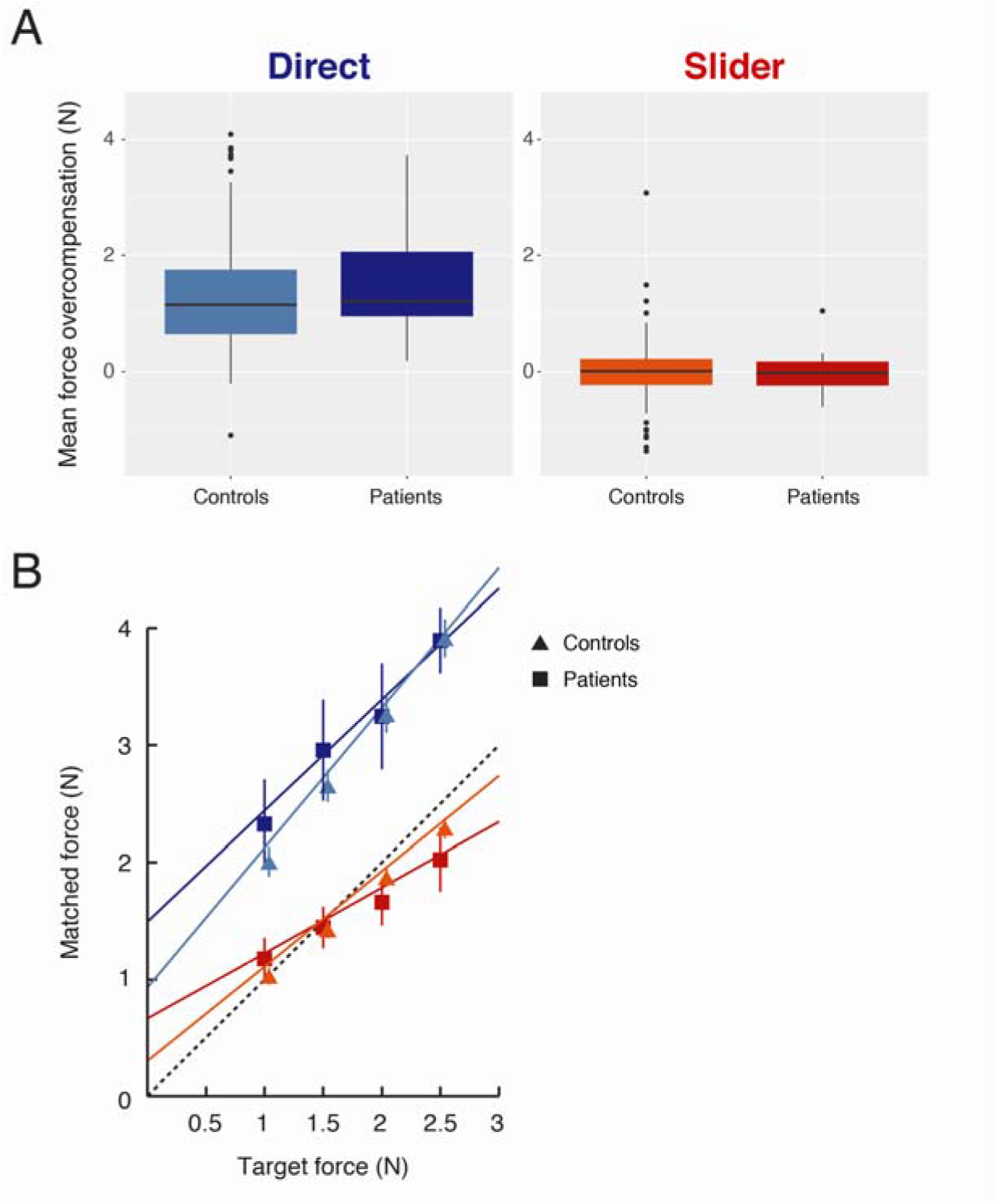
Differences in sensorimotor attenuation between PD patients and controls. A. Standard boxplots showing the distribution of mean force overcompensation values across all patients and controls in the Direct (shades of blue) and Slider (shades of red) conditions. Positive value indicates sensory attenuation. B. Mean regression plots of matched versus target force in the Direct and Slider conditions for both groups. Colour scheme is the same as in (A). Dashed line indicates the line of equality. Error bars indicate ±2 standard error of group mean. Control data points and error bars are offset by 2 pixels for illustration.

To examine changes in attenuation in PD, we compared overall attenuation and attenuation as a function of force levels in patients with our age-matched, normative control data. There were no significant differences in mean Direct force overcompensation (Mann-Whitney test; *Z* = 1.00, *p* = 0.32), suggesting no group difference in overall attenuation. Similarly, no group differences emerged in mean Slider overcompensation (*Z* = −0.45, *p* = 0.65). To compare attenuation as a function of force levels, we compared the intercept and slope of the linear regression fits of target force versus matched force. Specifically, changes in attenuation could be expressed as: (i) a shift in overcompensation across all forces, reflected in a different intercept with normal slope; and/or (ii) differences in sensitivity to the target forces, reflected in varying levels of overcompensation across force levels and changes in the slope.

We performed a linear regression of the matched force against target force for each condition and participant (Fig. 2B). The fit was better in controls, with patients showing smaller R^2^ compared to controls in both conditions (Direct: *Z* = 2.37, *p* < 0.05; Slider: *Z* = 3.30, *p* < 0.001). In the Direct condition, there was an increase in the intercept in patients compared to controls (Mann-Whitney test; *Z* = 2.25, *p* < 0.05, Bonferroni corrected), with no significant difference in the slope (*Z* = −1.8, *p* = 0.14, Bonferroni corrected). In the Slider condition, patients showed both reduced slope (Mann-Whitney test; *Z* = −2.82, *p* < 0.01, Bonferroni corrected) and a corresponding increase in intercept (*Z* = 2.75, *p* < 0.01, Bonferroni corrected), suggesting reduced sensitivity to the different forces. Together, these results resemble the effect of age on attenuation (Wolpe *et al*, 2016), with medicated PD patients showing consistent increase in attenuation of matched forces across the different force levels (despite no changes in overall attenuation here) but reduced sensitivity to externally generated forces.

Before testing how sensory attenuation might be related to clinical features in patients, we conducted several control analyses to verify the patients’ abilities to match the forces in the Direct condition. Compared to controls, patients showed increased within-trial variability in the forces they applied (Mann-Whitney test; *Z* = 2.72, *p* < 0.01), consistent with previous findings in PD (Stelmach *et al*, 1989). This is likely to arise from reduced force sensitivity due to increased sensory variability (Konczak *et al*, 2012), which, importantly, is not expected to introduce a systematic bias given the nature of the force matching task (see Discussion). However, increased within-trial force variability could also arise from the patients ‘overshooting’ and then slowly adjusting the force; an impaired ability to decide what force to apply; and/or from difficulties maintaining a steady force due to fatigue.

To explore the possibility of a *systematic* bias in the matching procedure, we performed additional analyses. First, for each trial, within the analysed time window, we fit a linear regression model of matched force against time. There was no linear trend in the matched force (regression slope not significantly different from zero; *t_17_* = −0.96, *p* = 0.35), similar to controls (Mann-Whitney test; *Z* = 0.53, *p* = 0.53). Moreover, there was no consistent relationship between this linear trend and the magnitude of the force applied by each patient (mean Pearson correlation coefficient not different from zero in patients; *t_17_* = −1.41, *p* = 0.18), which was again similar to controls (Mann-Whitney; *Z* = 0.15, *p* = 0.25). These results suggest it is unlikely that patients fatigued in the force they applied. Further, although not significantly different from controls, the slope in the Direct condition was overall smaller in patients relative to controls (see Figure 2B). This raises the possibility that patients were more limited in their ability to match larger forces. However, a closer look at the Direct slope demonstrated it was not significantly different from a veridical slope of one (Wilcoxon signed-rank test; *Z* = −1.76, *p* = 0.08). To further explore patient performance in matching larger forces, we computed the Pearson coefficient of the correlation between the magnitude of matched forces and within-trial SD for each trial and for each subject. There was a consistently positive relationship between the matched force and within-trial SD in patients (mean Pearson correlation coefficient significantly greater than zero across patients; *t_17_* = 8.09, *p* < 0.001), consistent with the well-known association between generated force variability and force level (Jones *et al*, 2002). Importantly, however, this association did not differ from controls (Mann-Whitney test; *Z* = −1.21, *p* = 0.23). Taken together, these control analyses suggest that PD did not alter the matching procedure itself. Nevertheless, we accounted for group differences in matching procedure (within-trial variability and R^2^ of the matched against target force) in the next analysis.

### Dopamine, disease severity and sensory attenuation

To examine the relationship between sensory attenuation and patient dopamine dose, we next fit a linear regression model with Direct force overcompensation as the dependent variable. The independent variables were disease severity, as assessed using the Unified Parkinson’s disease Rating Scale motor subscale III, and levodopa dose equivalency (LDE; Tomlinson *et al*, 2010). Additional variables that differed between groups were entered as covariates of no interest, including the Slider slope, within-trial force variability and the unexplained variance of the linear fits.

The regression model was statistically significant (*F*_(5,12)_ = 3.24, *p* < 0.05; 40% of force overcompensation variance explained; Fig. 3A). Even though disease severity and LDE were marginally positively correlated (*r_(16)_* = 0.46, *p* = 0.056), they had opposite effects on Direct overcompensation in patients. Disease severity was a negative predictor (*t_12_* = −2.62, *p* < 0.05), whereas LDE was a positive predictor (*t_12_* = 2.51, *p* < 0.05). For illustration, the direct relationship between attenuation and dopamine dose is plotted in Figure 3B. This pattern of results did not change when additionally co-varying for cognitive function in terms of Addenbrooke’s Cognitive Examination score; the laterality of dominantly affected side (*p* > 0.5 for the coefficients of both variables); or when entering Direct intercept (c.f. Wolpe *et al*, 2016) as the dependent variable. These results suggest that dopamine treatment could restore parkinsonism-related reduction in sensory attenuation.

**Figure 3.**
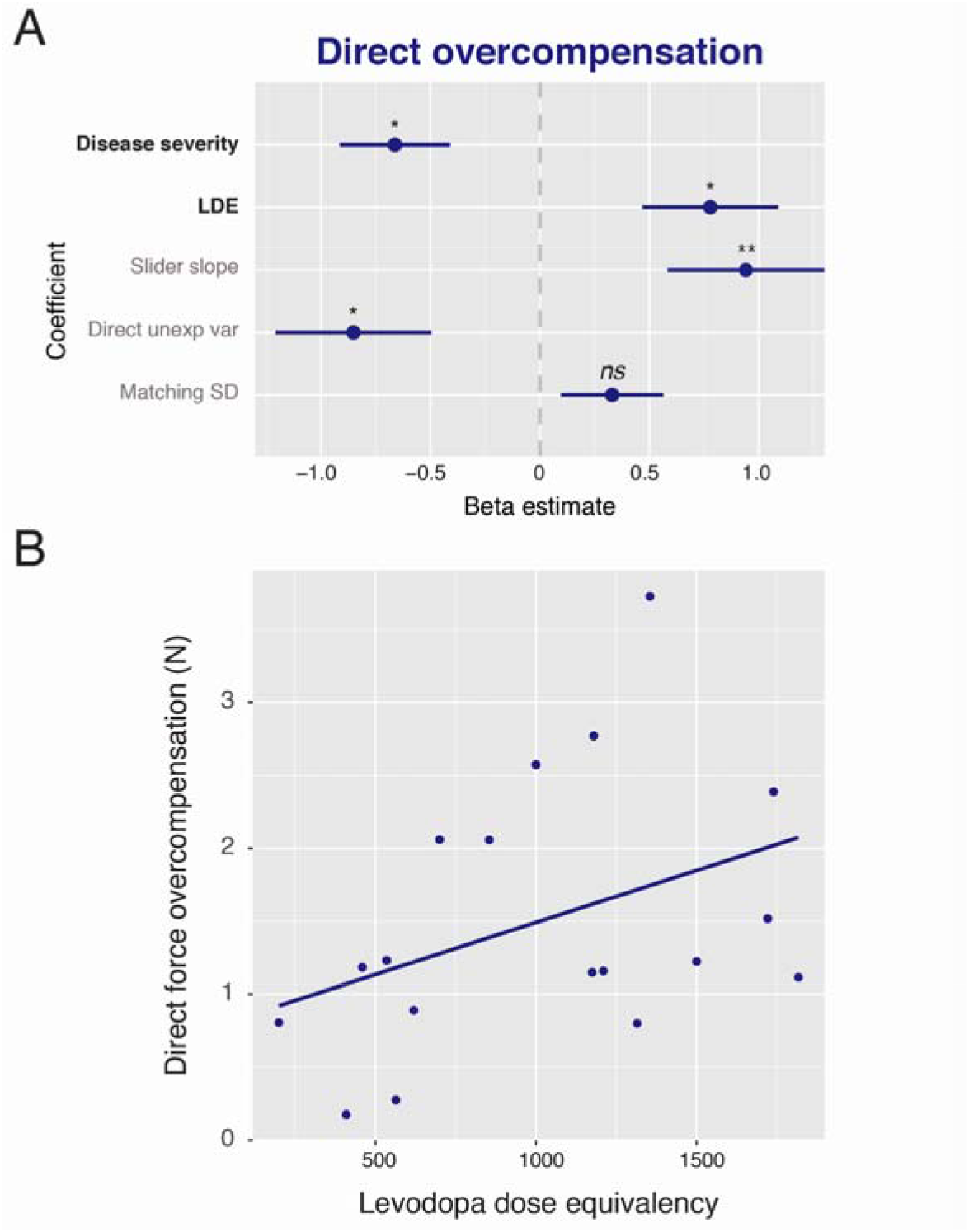
Association between dopamine and patient attenuation. A. Illustration of the standardised beta estimates of all independent variable coefficients included in the multiple regression model (R^2^_adj_ = 0.40), predicting Direct intercept. Clinical variables of interest were disease severity, which had a negative effect on attenuation, and levodopa doses, which had a positive effect on attenuation. Error bars indicate ±1 standard error of group mean. LDE = Levodopa Dose Equivalency. Significance level indicated by * = *P* < 0.05; ** = *P* < 0.01; *ns* = non-significant. B. Illustration of the relationship between Direct force overcompensation and levodopa dose equivalency, before entered into the regression model.

As Slider slope differed between groups, we fit a similar regression model to the Slider slope in a final exploratory analysis. Direct force overcompensation was included as a covariate of no interest, in order to identify predictors independently of attenuation effects included in the main regression model above. In this model (*F*_(3,14)_ = 6.82, *p* < 0.01; 51% of variance explained), disease severity was a positive predictor of Slider slope (*b* = 0.57, *t_14_* = 2.88, *p* < 0.05) and LDE was a significant negative predictor (*b* = −0.95, *t_14_* = −4.46, *p* < 0.001).

## DISCUSSION

The principal result of our study is that sensorimotor attenuation is positively related to dopamine doses in PD patients, whereas PD severity is related to reduced attenuation. These results can be interpreted in the context of optimal control theory, in which voluntary actions rely on the integration of sensory feedback with predictions of the consequences of one’s actions (Wolpert *et al*, 2011). The integration is precision-dependent, such that low-precision signals are down-weighted relative to high-precision signals. For example, this means that when performing a task in a dark or foggy setting, the precision of sensory feedback is expected to be low, and the sensorimotor system therefore relies more strongly on prior predictions when performing an action (Körding and Wolpert, 2004). Sensory attenuation is thought to reflect the precision of predictive signals (Bays *et al*, 2006), relative to the precision of sensory feedback (Brown *et al*, 2013; Wolpe *et al*, 2016). Our finding that PD motor severity is associated with reduced attenuation would therefore suggest that the precision of predictive signals may be compromised in PD (Macerollo *et al*, 2016). This may underlie the changes of sensorimotor integration in PD (Abbruzzese and Berardelli, 2003) and the dependence of patients on sensory cues, e.g. for the initiation and maintenance of their movement (Klockgether and Dichgans, 1994).

Although the severity of parkinsonism and doses of dopamine replacement therapy were positively correlated, dopamine was associated with an opposite effect, namely increased attenuation. These results support the hypothesis that dopamine alleviates disorders of movement in PD by restoring the precision and hence the typical reliance on sensorimotor predictions (Macerollo *et al*, 2016), at the expense of down-weighting the sensorium. This is further supported by the finding that an increase in dopamine dose was related to reduced force sensitivity in the Slider condition. These results are further consistent with the exaggeration of age-related sensory deficits in medicated PD patients (Konczak *et al*, 2012), and the detrimental effects of dopaminergic therapy on sensory sensitivity in PD (Konczak *et al*, 2009; O’Suilleabhain *et al*, 2001), although conflicting data on this exist (Li *et al*, 2010).

The relationship between dopamine and predictions has been previously tested indirectly in PD. For example, dopamine increases the perceived temporal attraction or ‘binding’ between an action and its effect (Moore *et al*, 2010). As binding critically relies on sensorimotor prediction (Moore and Haggard, 2008; Wolpe *et al*, 2013), these results are consistent with the hypothesis that dopamine increases the reliance on sensorimotor predictions. However, other studies reported the opposite association, in which dopaminergic treatment reduced the reliance on predictions in perceptual decision-making tasks, while increasing reliance on sensory information (Vilares and Kording, 2017; Wolpe *et al*, 2015). We propose that these are different types of predictions that are mediated by distinct brain mechanisms within a cortical hierarchy (c.f. Brown *et al*, 2013). “Low-level” sensorimotor predictions reflected in sensory attenuation depend on pre-SMA connectivity with dorsal striatum circuits (Wolpe *et al*, 2016), and could play a key role in normal execution of movement (Brown *et al*, 2013; Macerollo *et al*, 2016). On the other hand, high-level perceptual priors depend more on prefrontal connections with ventral striatum circuitry (Vilares *et al*, 2012; Wolpe *et al*, 2014). Since dopamine doses are tailored to alleviate patient motor symptoms, which mostly reflect dorsal striatal dopamine depletion, high dopamine doses can effectively “overdose” the ventral striatum (Cools, 2006). This discrepancy might lead to the relative normalisation of low-level predictions for attenuation, but a weakening of high-level predictions for perceptual decision making tasks.

The positive association between attenuation and dopamine, and the combination of increased Direct intercept and reduced Slider slope in medicated PD patients mirror the impact of healthy ageing on sensory attenuation (Wolpe *et al*, 2016). Increased attenuation found with normal ageing is associated with reduced connectivity in a fronto-striatal network (Wolpe *et al*, 2016) that is strongly affected in PD. This network includes the caudate and putamen, dorsolateral prefrontal cortex and pre-SMA as the network hub. Interestingly, reduced fronto-striatal connectivity has also been associated with increased caudate dopamine synthesis as seen in healthy ageing (Klostermann *et al*, 2012). Therefore, increasing dopamine synthesis – including by levodopa administration in PD – could increase sensory attenuation by altering connectivity of the pre-SMA within its fronto-striatal network.

Activity in the secondary somatosensory cortex, mediated via increased pre-SMA connectivity (Shergill *et al*, 2013; Wolpe *et al*, 2016), has also been suggested as the neural correlates of attenuation. Neurophysiological studies have indeed demonstrated that the perceived attenuation is closely related to *late* components of sensory evoked potentials, arising from the secondary somatosensory cortex (Palmer *et al*, 2016). In PD, however, *early* components of sensory evoked potentials, arising from fronto-striatal activity, are already altered (Abbruzzese and Berardelli, 2003). The typical neurophysiological attenuation of these early components following a voluntary movement is absent in PD patients ‘off’ medication, and restored by dopamine treatment (Macerollo *et al*, 2016). This reduced neurophysiological attenuation of the early components of sensory evoked potential has been attributed to a failure in sensory ‘gating’ in PD (Macerollo *et al*, 2016), resulting from abnormal precision of sensory afferents (Brown *et al*, 2013).

The neurophysiological gating of sensory afferent signals before and during movement has been proposed to be required for the initiation processes of voluntary movements (Barker, 1988; Brown *et al*, 2013). This theoretical account suggests that a relative reduction in the precision of sensory signals, leading to sensory attenuation, enables high-level predictions to drive normal movement through hierarchical networks in the central nervous system (Brown *et al*, 2013). In PD, deficient precision of predictions would be overwhelmed by sensory evidence for a lack of movement, resulting in bradykinesia (Brown *et al*, 2013). Although the neurophysiological attenuation of early components of sensory evoked potentials, shown to be impaired in PD and remediated with dopamine, may not directly underlie the behavioural phenomenon of attenuation (Palmer *et al*, 2016), our behavioural results are consistent with these previous studies. Our findings that PD motor severity is associated with reduced attenuation while dopamine dose is related to increased attenuation, support the hypothesis that bradykinesia in Parkinson’s disease could be considered in terms of pathological imprecision of sensorimotor prediction, which are alleviated by dopamine treatment.

Our results have interpretative limitations that should be considered. Firstly, we opted for the force matching task, as it is a simple, highly intuitive and robust task (Wolpe *et al*, 2016). Importantly, in a matching task any absolute bias is factored out and therefore sensory deficits, which are common in PD (Konczak *et al*, 2012), can increase performance variability, as observed in our study, but are unlikely to introduce a systematic bias (e.g. with a force of ‘X’ N sensed as X/2 N, patients would still have to match X N to experience the same X/2 N intensity). However, the task does have its limitations and preliminary testing showed that the it is not easily performed by PD patients ‘off’ medication, mainly because of tremor and akinesia in light of the timed, dexterous movements required for matching the forces. Patients were thus tested in an ‘on’ state, to avoid the confound of severe performance problems, and we draw inferences from the between-subject variability in disease severity and dopamine doses. We therefore cannot draw causal inferences on the effect of dopamine in patients. Secondly, the reduced slope in the Direct and Slider conditions suggests that reduced sensory sensitivity may have dominated patient behaviour in the task (Konczak *et al*, 2012). Importantly, differences in patient force sensitivities were accounted for in main regression analyses, together with other potential confounders, suggesting that reduced force sensitivity was not a sufficient explanation for the significant associations between attenuation, disease severity and dopamine dose.

In conclusion, our study suggests that dopamine is related to an increase in sensory attenuation in PD, suggesting that dopamine increases the precision of sensorimotor predictions. The results support the hypothesis that bradykinesia in movement disorders like PD can, in part, be considered in terms of pathological (im)precision of sensorimotor predictions (Brown *et al*, 2013), which can be modulated by dopamine (Macerollo *et al*, 2016). This may provide a common framework for understanding the role of dopamine in perceptual, cognitive and motor function.

## ACKNOWLEDGEMENTS

We are grateful to the patients for their participation in the study. We are also indebted to the Cam-CAN respondents and their primary care teams in Cambridge for their participation. We thank Professor Roger Barker for his helpful comments on the manuscript and the support of the John van Geest Centre for Brain Repair, Parkinson’s disease research clinic.

